# Trap Spacing Protocols for Amazonian Dung Beetle Surveys

**DOI:** 10.1101/2024.07.16.603696

**Authors:** Elena Hernandez Perez, Pablo Aycart, Elena Chaboteaux, Alejandro Lopera Toro, Adrian Forsyth

## Abstract

Surveys of dung beetle species richness, abundance and assemblage composition were conducted at Los Amigos Biological Station in Amazonian Peru, using dung-baited pitfall traps with varying spacing and duration. Trap spacing did not significantly influence species richness, the number of individuals captured, or assemblage composition. These findings suggest that in healthy Neotropical rainforests, dung beetle populations are vast and mobile, indicating that the concern over trap spacing might be less critical than previously considered.

## 1. INTRODUCTION

Dung beetles are a conspicuous and ecologically important component of Neotropical forests (Peck & Forsyth, 1982; Bicknell et al., 2014, Viegas et al., 2014, Cajaiba et al., 2018). They play important roles in secondary, and less often primary, seed dispersal (Andresen, 2002; Vulinec, 2002; Whitworth, 2021; Chaboteaux et al., 2023), bioturbation (Nichols et al., 2008; Milotic et al., 2017), nutrient recycling (Maldonado et al. 2019; Noriega et al. 2007), fecal parasite destruction (Sands & Walls, 2017) and display a high species diversity and functional richness (Slade, 2007). Furthermore, they exhibit complex behaviors related to competition for and consumption of dung, carrion, fungal, detrital and other food resources (Simmons & Ridsdill-Smith, 2011). Dung beetles rely heavily on mammalian excrement (Noriega, 2012) for sustenance and reproduction (Nichols et al., 2008), making them particularly vulnerable to environmental changes. This underscores the critical importance of preserving both dung beetles’ and vertebrates’ intricate communities and their ecosystem’s relationships (Spector, 2006). Dung beetles could serve as effective indicators of ecological health resilience (Whitworth et al., 2021; Noriega et al., 2020), providing insights into the success of conservation strategies over time.

Most assemblages show changes in response to disturbances such as deforestation and selective logging (França et al., 2017; Slade et al., 2011), fragmentation (Nichols et al., 2007; Andresen, 2003), introduction of livestock (Guerra Alonso et al., 2022), hunting pressure on mammals (Nichols et al., 2009), and a sensitivity to thermal alterations associated with climate change (Menéndez et al., 2014; Gaston & Chown, 1999).

Most species are readily and rapidly collected with baited pitfall traps (Aycart et al., 2023; Mora-Aguilar et al., 2023), making them a commonly utilized ecological indicator (Noriega et al. 2020; Davis et al., 2001; Bicknell et al., 2014) and an essential component in biodiversity surveys. This has led to proposals for standardized sampling protocols in order to obtain comparable data sets for different times and places (Mora-Aguilar et al., 2023). Attention is being paid to trap spacing based on the premise that spatially independent traps will avoid pseudo-replication and ensure more efficient surveys. Commonly recommended and utilized spacings are often from 50 m (Larsen & Forsyth, 2005) to 100 m (Da Silva & Hernandez, 2015), but there are suggestions that even greater separation might be needed (Ong et al., 2022). These proposed spacings are primarily based on mark and recapture data sets that measure how far a dung beetle might fly in a limited time period.

There are several reasons why trap spacing would be important when surveying dung beetle species’ diversity and abundance. If traps are located too close to one another, they may capture a similar and limited subset of the dung beetle community (Marsh et al., 2013). The assemblages found in a localized area are likely to be more similar in terms of species composition due to similar environmental conditions and limited movement over short distances (Judas et al., 2002; Gibb et al., 2006). As a result, surveys using closely placed traps might not sample assemblages as effectively as wider-spaced traps (Ward et al., 2001). Moreover, varying microclimatic conditions may support different species of beetles (Grimbacher et al., 2006; Lucio-García et al., 2022). If the sampling is concentrated in a small area with high spatial autocorrelation, it will not capture the heterogeneity of the area under study (Blanchet et al., 2013; Moctezuma, 2021).

However, against these considerations, researchers must weigh the time and cost requirements of wide trap spacing and the effect of transect disturbance on vegetation when an established trail system is not being used for trap placement. In addition, these trap spacing may not be feasible in steep terrain or islands and forest fragments (Larsen et al., 2008).

Accordingly, we conducted a variety of trials, altering trap spacing to study how estimates of dung beetle species richness, abundance and community composition vary depending on the distance between pitfall traps within transects in an Amazonian lowland rainforest.

## 2. METHODS

### 2.1 Study site

Los Amigos Biological Station is located in the Madre de Dios region, in southeastern Peru on a private property of 453 ha that is contiguous with a conservation concession totaling 145,918 ha (12°34’07”S 70°05’57” W; 225–296 m a.s.l.). The concession touches the eastern flank of Manu National Park, acting as a corridor for wildlife, and the southern borders of the Reserva Territorial Madre de Dios. Both the station and concession are part of a vast wilderness with the complete megafauna of a dozen primate species and abundant terrestrial mammals, which have been protected against hunting and logging since the establishment of the concession in 2001. The region has a dry season from May to September and a wet season from October to April. The study site presents both old-growth and secondary forests, extending across high terrace (*terra firme*) and floodplain habitats (Martel & Cairampoma, 2012), with soils mainly classified into ultisol and inceptisol orders (Nikitina et al., 2011). Sampling was restricted to mature forests with minimal anthropogenic disturbance. The dung beetle fauna of Los Amigos has been the subject of other extensive dung beetle surveys in the past (Larsen et al., 2008; Aycart et al., 2023).

### 2.2 Study design

Dung beetles were captured using human dung-baited pitfall traps. The traps consisted of 16 oz (473 ml) plastic cups half-filled with water, detergent and salt, which were placed in the ground. The bait (30 g) was wrapped in a gauze bandage tissue which was tied to a stick so that it hung a few centimeters above the cup. Each trap was covered by a polystyrene plate to prevent the trap from being inundated by rain and to stop beetles from landing directly on top of the bait (Martínez-Hernández et al., 2022). Subsequently, dung beetles were identified to species level using Larsen and Génier (2008), Vaz-de-Mello et al. (2011), Figueroa et al. (2014), Silva and Valois (2019), and Rossini and Vaz-de-Mello (2020). GPS locations were recorded for each trap location to check for spatial autocorrelation. Sampling was conducted equally in *terra firme* and floodplain habitats.

Two different trials to assess trap spacing effects were conducted:

a. Trial A. Traps were deployed in four transects of five traps per transect with different inter-trap spacing within a transect: 1 m, 5 m, 50 m and 100 m. Traps were sampled every 48 hours and after this time, traps were collected and the transects were rotated between trails. Two repetitions were run during the rainy season. The first repetition was in October 2021 and the second was in January 2022. A total of 160 pitfall trap samples were deployed; however, 2 traps were lost due to animals. Hence, a total of 158 trap samples were included in the analysis.
b. Trial B. A binary comparison was made between traps spaced 5 m apart and those spaced 50 m apart. Traps were deployed in two transects of two traps with the same trap distance. Similar to trial A, traps were collected every 48 hours and the transects rotated between trails, hence using both trails for the same transect. This experiment was run in April and May 2023 and a total of 56 pitfall traps were collected.

This study did not consider other factors that might influence dung beetle capture rates, habitat heterogeneity, and seasonal changes (Pessoa, 2021). Furthermore, this research was confined to a specific location, which may limit the generalizability of our findings.

### 2.3 Statistical Analysis

All statistical analyses were conducted using R software (R Core Team, 2023).

A generalized linear mixed model (GLMM) was used to investigate: a) the trap-level alpha diversity, comparing the number of species (species richness) and individuals (abundance) found in each trap depending on the distance to the nearest trap; b) the transect-level alpha diversity, comparing the species richness and abundance found in each transect for each trap spacing within a particular transect. This approach quantifies the extent to which these ecological variables are influenced by the spatial separation of the traps for trials A and B. The package glmmTMB (Brooks et al., 2017) was employed. A Poisson distribution was first selected due to the nature of the data, but in case of overdispersion, a non-binary distribution was employed. The models included trap distance as a fixed effect, with habitat, trail, and repetition (this last one only in trial A) considered as random effects to control for variability unattributed to trap distance. To study residuals, the DHARMa package (Hartig, 2018) was employed here and every time a GLMM model was run throughout the analysis. Additionally, ANOVA was performed using the Anova function from the car package (Fox & Weisberg, 2018) to compare the effects of the fixed factors in the model.

To understand the effect of trap distance on the number of species detected from the entire pool of all sampled species, the total number of detected species was compared across all traps for each treatment (trap distance). Additionally, a sample coverage curve was produced for all distances using the iNEXT package (Chao et al., 2014). The data for each assemblage was prepared and the iNEXT function was run, specifying the abundance-based datatype. This produced species accumulation curves, which were used to evaluate the completeness of the sampling effort across different distances.

Beta diversity, considered as species turnover, was quantified at the transect-level using the Soerensen index, a measure of species similarity, while taking into account species abundances. For this, BAT package (Cardoso & Carvalho, 2015) was employed. A GLMM with a Gaussian family was then applied to examine the relationship between beta diversity and trap distance.

The model included random effects for both habitat and trail to account for the non-independence of samples within these categories. When encountering the non-normality of the data in trial A, this was inversely transformed in order to stabilize the variance and approximate a normal distribution.

To assess the composition and distribution of species within the study area, two key diversity indices were calculated: the Shannon index and the Inverse Simpson index. The Shannon index is a measure of species diversity in a community, taking into account both species richness (the total number of species) and evenness (the relative abundance of each species). The Inverse Simpson index is another measure of diversity, which gives more weight to the more abundant species in the community. It is the reciprocal of the Simpson index, which is the probability that two randomly selected individuals from the sample belong to the same species. These metrics provided insights into the diversity and distributional equity of species assemblages. The package iNEXT (Chao et al., 2014) was employed for this purpose. Data was grouped by transect-level, so the indices were calculated for the same trap distance a total of eight times, corresponding to the amount of times that transects were rotated.

A GLMM was employed to analyze the impact of trap distance on biodiversity, specifically focusing on the Shannon and Simpson diversity indices, with trap distance as the predictor variable, and assuming a Gaussian distribution due to the continuous nature of the response variables.

## 3. RESULTS

In trial A, the total amount of species recorded was 91, and the total abundance of individuals was 17,623. The most common species were *Onthophagus osculatii* Guérin-Méneville, with a total count of 4,361; a morphospecies of *Onthophagus* Latreille that could not be identified to species level and was labeled as *Onthophagus morphospecies 1*, with a total count of 1,506; and *Onthophagus transisthmius* Howden & Young, with a total count of 1,503.

On the other hand, in trial B, there were 108 species recorded and the total abundance of all species combined was 8,150. The three most common species found were *Eurysternus caribaeus* (Herbst) with an abundance of 1,113; *Eurysternus vastiorum* (Martinez) with a total count of 1,035; and *Sylvicanthon proseni* Martinez with an abundance of 612.

The combined trials contained 154 species in total and the combined total abundance of all species was 25,773. The three most abundant species across both experiments were *O. osculatii* with an abundance of 4,361; followed by *E. vastiorum* with an abundance of 2,208 and *Onthophagus morphospecies 1* with an abundance of 1,506.

### 3.1 Trap-level alpha diversity

For trial A, the species richness in each trap depending on the distance to the nearest trap was studied. The results indicated no significant effect of trap distance (at 1, 5, 50, and 100 meters) on dung beetle species richness (p-values: 0.4872 > 0.05). Hence, within the distances studied, the spatial arrangement of traps does not significantly affect the capture of species richness. Random effects analysis showed the presence of variability in species richness, attributed to habitat, trail, and repetition, with the repetition random effect exhibiting the highest variability, but they were non-significant. Finally, to evaluate model fit, overdispersion was tested and found to be non-significant, indicating that the Poisson model was appropriate for the data. Residual evaluations confirmed that the residuals were in accordance with the expected distribution, showing no significant deviation in normality nor in dispersion or outlier presence.

The same procedure was done for the variable abundance, which yielded no significant effect of trap distance on the number of individuals (p-values: 0.4869 > 0.05). Again, the repetition random effect showed the highest variability. No overdispersion was detected and residuals did not reveal any significant deviations from the expected distribution.

For trial B, the analysis showed no significant effect of trap distance on species richness (p-value = 0.7134 > 0.05). This implies that the proximity of traps does not substantially affect the detection of different dung beetle species. Random effects were small, with variance components indicating a slight influence of habitat and trail on species richness, though neither was statistically significant. Residual diagnostics did not reveal any issues with overdispersion, and plots of residuals versus predicted values did not reveal any substantive issues.

The same method was used for the variable abundance, which showed no significant impact of trap distance on the number of individuals (p-values: 0.4683 > 0.05). No overdispersion was observed, and the residuals did not indicate any significant deviations from the expected pattern.

### 3.2 Transect-level alpha diversity

Species richness for each transect was studied to check if the distance between traps affects the number of species found in each transect. For trial A, results from the models indicate that trap distance did not have a significant effect on species richness (p-value = 0.827 > 0.05). This suggests that the different distances at which traps were placed in each transect did not affect the total number of species captured. Random effects analysis pointed to minimal variability attributed to the habitat and trail, with the repetition effect showing a slightly higher but still small variance. Residual checks showed no significant overdispersion and the residuals plotted against predicted values showed no significant problems, indicating that the model assumptions were not violated. In trial B, the analysis demonstrated that the distance between traps, at 5 and 50 meters, did not have a statistically significant effect on the richness of dung beetle species (p-value = 0.1176). This suggests that the species richness is not influenced by the spacing of traps within each transect, indicating a uniform spatial distribution of dung beetle species richness. The random effects indicated minimal variation due to habitat and a slightly higher variation due to trail, but neither random effect was significant enough to affect estimates. The model’s adequacy was further supported by diagnostic checks, which showed no evidence of overdispersion. Additionally, residual diagnostics performed did not reveal any significant deviations from the expected distribution.

Abundance for each transect was studied to check if the distance between traps affects the number of individuals found in each transect. For trial A, the results demonstrate that trap spacing distance did not significantly affect the abundance of dung beetles (p-values: 0.9944 > 0.05). This implies that the number of dung beetles captured is consistent across transects at varying distances, indicating a uniform distribution of individuals within the study area. The model showed minimal random effect variance for habitat and trail, while the variance for repetition was somewhat larger, though it did not contribute significantly to the model. Residual checks confirmed no significant overdispersion, and plots of residuals versus predicted values did not reveal any substantive issues. In trial B, findings revealed no significant effect of trap distance on the abundance of dung beetles (p-value = 0.1581 > 0.05). This result suggests that the number of individuals captured is not affected by the varying distances (5 and 50 meters) between transects. The random effects analysis revealed some variation attributed to the habitat and a greater variation due to the trail; however, neither contributed significantly to the overall abundance of species captured. The model’s adequacy was confirmed through checks for overdispersion, with no issues detected. Additionally, the diagnostic plots showed residuals that were consistent with the model’s assumptions, exhibiting no significant deviations from expected patterns.

### 3.3 Total species richness

In trial A, for each of the trap spacing (1, 5, 50 and 100 m), the sample coverage curves indicate that as sampling continues, the curves approach an asymptote, suggesting that additional sampling leads to diminishing returns in terms of discovering new species (Figure 1). The curves for distances are very close to each other, indicating that the sample coverage is similar at all distances. This suggests that sampling at 1, 5, 50 and 100 meters yields comparable representations of the community. The curves indicate that around 80% of the community is covered when about 40 samples are taken at 1, 5, 50 and 100 meters. This suggests that the sampling effort is relatively complete at these distances, as additional sampling would only yield marginal increases in sample coverage.

**Figure 1.**
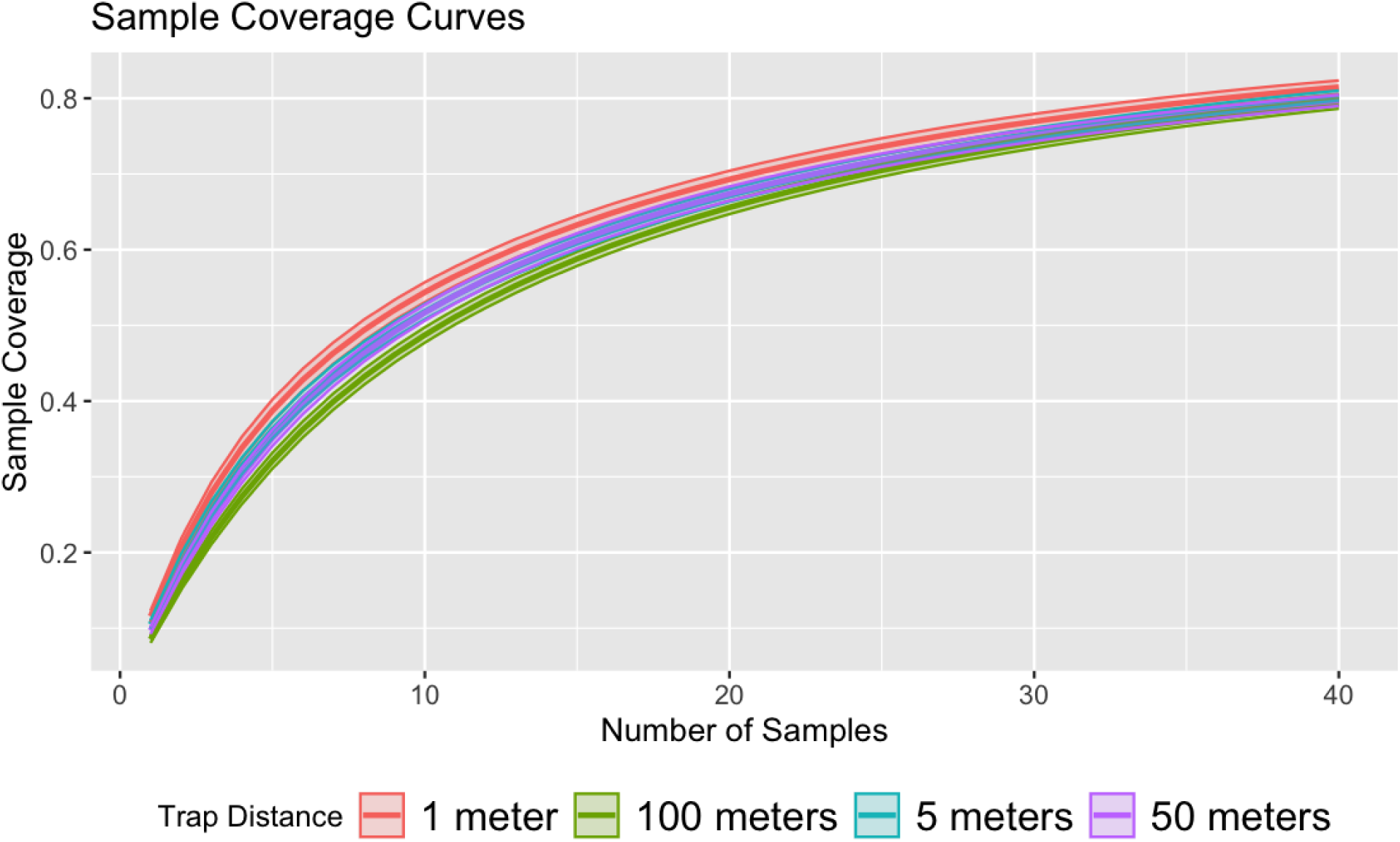
Sample coverage curves for the 1 (red line), 5 (blue line), 50 (purple line) and 100 m (green line) distances show the relationship between the number of samples and the proportion of the total community represented in those samples.

Similarly, in trial B, the sample coverage curves plotted from the data derived from the two trap distances, 5 and 50 m, (Figure 2) show a similar pattern to that observed in trial A, overlapping even more.

**Figure 2.**
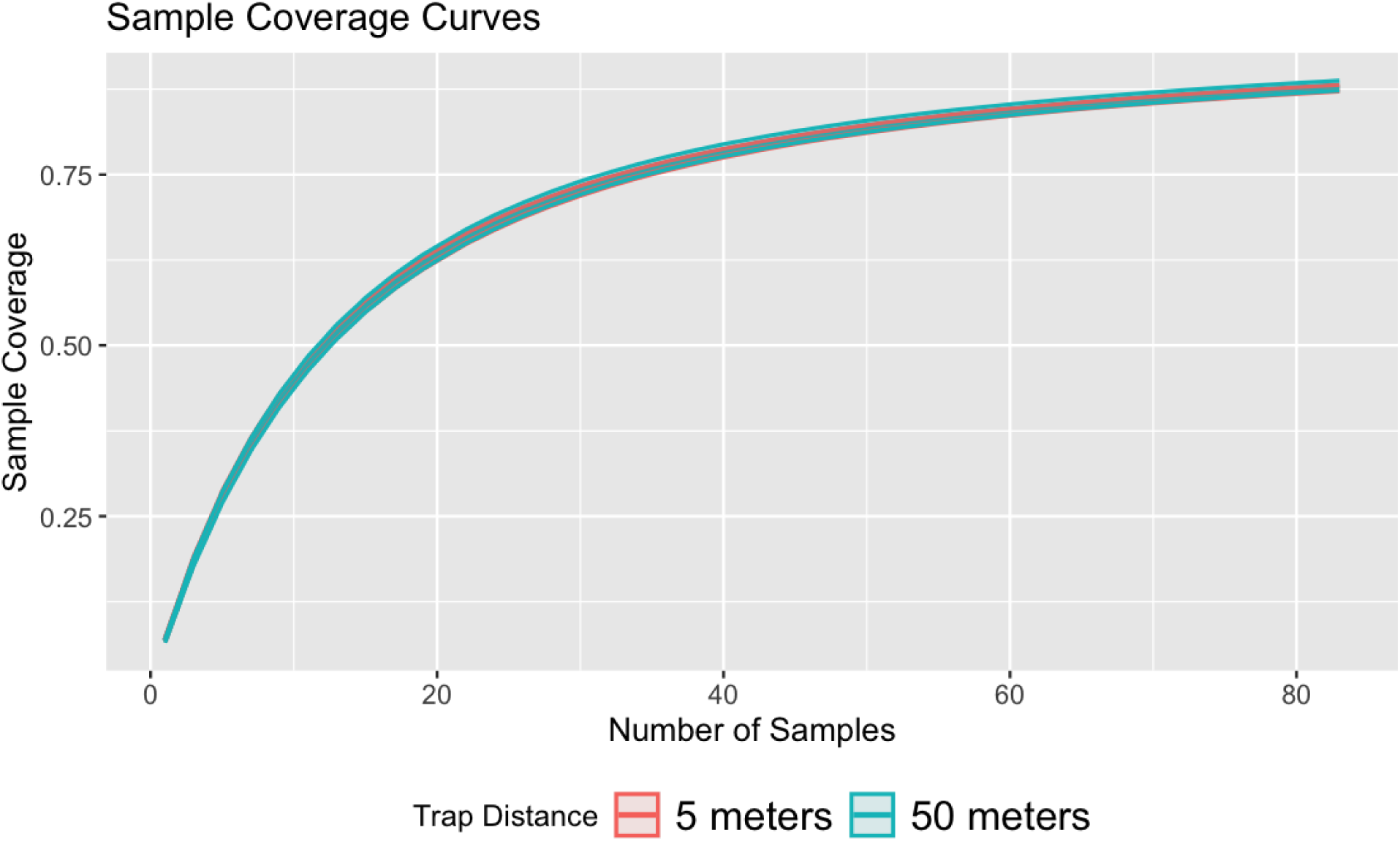
Sample coverage curves for the 5 (red line) and 50 m (blue line) distances show the relationship between the number of samples and the proportion of the total community represented in those samples.

### 3.4 Transect-level beta diversity

In trial A, the fitted GLMM revealed a conditional model with a non-significant effect of trap distance on beta diversity (p-value = 0.0947 > 0.05), suggesting that trap spacing did not significantly influence species turnover when controlling for habitat and transect effects. Model diagnostics did not indicate the presence of overdispersion. The residual analysis suggested an adequate fit of the GLMM to the data.

In trial B, the GLMM analysis also yielded a conditional model in which trap distance had no statistically significant impact on beta diversity (p-value = 0.193 > 0.05), indicating that the spacing between traps was not a significant factor in species turnover after accounting for the effects of habitat and transect. Additionally, diagnostic checks revealed no overdispersion, and the residuals suggested that the GLMM provided a satisfactory representation of the data.

### 3.5 Diversity indices

In trial A, for the Shannon diversity index, the GLMM analysis revealed no significant effect of trap distance (p-value = 0.629 > 0.05). Diagnostic checks for overdispersion confirmed that the model’s dispersion was appropriate and no overdispersion was detected. Residual diagnostics displayed a homogeneous spread of residuals against predicted values with no significant deviations from expected.

Similarly, the model for the Simpson diversity index did not display a significant effect of trap distance (p-value = 0.509 > 0.05). No overdispersion was present in the model. The residual diagnostics demonstrated an adequate fit, with residuals evenly scattered across the range of predictions and conforming to the expected normal and no significant dispersion or outlier issues.

In trial B, the Shannon index model outputs demonstrated that trap distance had no statistically significant influence (p-value = 0.722 >0.05). The diagnostic procedures conducted did not detect any overdispersion. Residual evaluations confirmed that the residuals were in accordance with the expected distribution, showing no significant deviation in normality nor in dispersion or outlier presence.

Concerning the Simpson index, similar conclusions were drawn with no discernible impact of trap distance (p-value = 0.817). Model validation checks revealed no overdispersion, and residual diagnostics validated the model fit, indicating residuals that conformed to the expected normal distribution without notable dispersion or outliers.

## 4. DISCUSSION

One of the first attempts at estimating dung beetles’ abundance in a Neotropical rainforest (Peck & Forsyth, 1982) found a low recapture rate implying a high population size and/or high mobility. The trap spacing experimental results reported here support this notion: dung beetle species distribution and abundance are robust to variations in the spatial arrangement of sampling traps within the measured distances. Wide trap spacing of 100 m or more as a mean to achieve spatial independence between traps may not be necessary if the intent of a baited pitfall survey is to rapidly assess dung beetle species richness and abundance in healthy Amazonian forests.

Earlier recommendations by Larsen and Forsyth (2005), and Da Silva and Hernandez (2015) who advocated for increased spacing to avoid sample overlap were based on the notion that traps have an “effective sampling radius” (ESR) that attracts beetles from an area defined by this radius of attraction. Here, it is suggested that the notion of using the ESR concept may not apply due to the fact that dung beetles in an intact Amazonian rainforest are abundant and highly mobile. Moreover, if the intent of a survey is to detect functional rarity, or merely rarity itself (Aycart et al., 2023), investing in diverse sampling methods and extending the duration of sampling may yield greater benefits than focusing solely on trap spacing.

Furthermore, utilizing reduced trap spacing may help minimize negative effects that the opening of new trails may have on ecosystem processes (Comita & Goldsmith, 2008). It is encouraged to any investigator developing a study of dung beetle species richness, abundance and composition to carefully consider all the various factors that will affect the time required and the cost of a survey, such as trap spacing. Moreover, consideration of seasonality (Andrade et al., 2011), microhabitat (Mehrabi et al., 2014), heterogeneity (Rivera et al., 2020) and bait type (Marsh et al., 2013) should all be contemplated.

Finally, these results give insights into the nature of dung beetle assemblages in a pristine Amazonian rainforest. The use of baited pitfalls is akin to trying to empty the ocean with a cup, given the high abundance and mobility of dung beetles. More detailed estimates of dung beetle population density and dispersion are needed to further refine dung beetle sampling protocols.

## 5. CONCLUSION

This study at Los Amigos Biological Station in Peru explored how trap spacing affects dung beetles’ diversity metrics like species richness and abundance in Amazonian assemblages. Trap spacing did not significantly affect the measurements of species richness, abundance or species assemblages composition. We conclude that those seeking to survey these characteristics in an Amazonian rainforest should re-evaluate the recommended trap spacing distances suggested by Larsen and Forsyth (2005), Da Silva and Hernandez (2015), or Ong et al. (2022). Overall, our results suggest that the “effective sampling radius” concept may not be the best paradigm for determining trap spacing in dung beetle surveys.

## ACKNOWLEDGEMENTS

We would like to express our gratitude to the Andes Amazon Fund (AAF), the International Conservation Fund of Canada (ICFC), and the Gordon and Betty Moore Foundation (GBMF) for their financial support and backing of this research. Our thanks also go to the Asociación para la Conservación de la Cuenca Amazónica (ACCA) for hosting us at the Los Amigos Biological Station (LABS) and Manu Biological Station (MBS). The beetles were collected under the SERFOR permit AUTIFS-2021-021.

## CRediT authorship contribution statement

**Elena Hernandez Perez**: Formal Analysis, Investigation, Methodology, Data curation, Writing – original draft. **Pablo Aycart**: Formal Analysis, Investigation, Methodology, Data curation, Writing – review & editing. **Elena Chaboteaux**: Investigation, Methodology, Data curation, Writing – review & editing. **Alejandro Lopera Toro**: Investigation, Methodology, Data curation, Writing – review & editing. **Adrian Forsyth**: Conceptualization, Funding acquisition, Investigation, Methodology, Writing – review & editing.

